# Host and environmental determinants of microbial community structure in the marine phyllosphere

**DOI:** 10.1101/2020.05.07.082826

**Authors:** Margaret A. Vogel, Olivia U. Mason, Thomas E. Miller

## Abstract

Although seagrasses are economically and ecologically critical species, little is known about their blade surface microbial communities and how these communities relate to the plant host. To determine microbial community composition and diversity on seagrass blade surfaces and in the surrounding seawater,16S rRNA gene sequencing (iTag) was used for samples collected at five sites along a gradient of freshwater input in the northern Gulf of Mexico on three separate sampling dates. Additionally, seagrass surveys were performed and environmental parameters were measured to characterize host characteristics and the abiotic conditions at each site. Results showed that *Thalassia testudinum* (turtle grass) blades hosted unique microbial communities that were distinct in composition and diversity from the water column. Additionally, results suggested that environmental conditions, including water depth, salinity, and temperature, were the major driver of community structure as blade surface microbial communities varied among sites and over sampling dates. Host condition may be a secondary driver of community structure as compositional changes were also correlated with host characteristics, including leaf growth rates and blade nutrient composition, Additionally, 21 microorganisms from five phyla (Cyanobacteria, Proteobacteria, Planctomycetes, Chloroflexi, and Bacteroidetes) were present in all blade surface samples and may represent a core community for *T. testudinum*. Members of this core community may have ecological importance for determining community structure or in performing key community functions. This study provides new insights and understanding of the processes that influence the structure of marine phyllosphere communities, how these microbial communities relate to their host, and their role as a part of the seagrass holobiont, which is an important contribution given the current decline of seagrass coverage worldwide.

## Introduction

In recent years, there has been an increasing number of studies on the microbial communities associated with the leaf surfaces, or phyllosphere, of terrestrial plants [1–5]. We now know that the phyllosphere is a rich habitat for microbes and that leaf surfaces can host up to 10^7^ microbial communities per cm^2^ of tissue. The composition and structure of phyllosphere microbial communities has been found to be driven by both environmental conditions, including precipitation/moisture and temperature [3, 6–10] and host plant identity [11, 7, 8]. Additionally, these leaf surface microbial communities can have a variety of relationships with their host plants ranging from beneficial to pathogenic [3]. Of particular interest is when microbial communities can act as a form of defense for the plant host by inhibiting invasion and/or colonization of potential plant pathogens as well as promote plant growth [1, 12]. In this way, these microbial communities can have positive interactions with their hosts which can ultimately affect host fitness and performance [4, 13] and create feedbacks that lead to the maintenance of a healthy phyllosphere microbiome. However, these beneficial host-microbe relationships can be transient due to changes in environmental or host condition.

In comparison to the terrestrial phyllosphere, much less is known about the microbial communities associated with the leaf surfaces of aquatic plants. Unlike terrestrial leaf surfaces, where microbial community members must withstand extreme changes in moisture, the marine phyllosphere is continually submerged in water, removing desiccation as an abiotic stressor and structural determinant. As in the terrestrial phyllosphere, these submerged leaf surfaces are sites for bacteria colonization and the formation of biofilms [14, 15, 16]. Both environmental conditions and the plant host can influence the micro-environment that the leaf surface microbial community experiences [5]. However, the biofilm itself has the potential to be a buffer to the conditions in the surrounding water column. This can lead to more stable microbial community dynamics over time than is seen on typical aerial leaf surfaces where abundance often depends on moisture availability [1, 6]. Due to this, the influence of the plant-host (through nutrient exudation and secondary metabolite production) may be more important for determining microbial community structure in the marine environment relative to the environmental conditions of the water column. This could create stronger plant-microbe interactions and different dynamics than are found in the terrestrial phyllosphere. However, the leaf-associated microbial communities of aquatic plants and the relationships these communities have with their plant-host remain largely unexplored.

Seagrasses are economically and ecologically critical species that occur in coastal waters worldwide [17, 18]. These marine angiosperms can act as ecosystem engineers forming dense meadows that serve as nurseries, feeding grounds, and habitats for a wide variety of marine species from tiny invertebrates to sea turtles and manatees. As foundation species, seagrasses support microbes, algal epiphytes, and their grazers creating highly productive ecosystems [17, 19]. In addition, seagrasses provide valuable ecosystem services, such as stabilizing sediments and trapping and cycling nutrients [20–22], and are important sites for blue carbon sequestration [23, 24]. However, with increasing anthropogenic influence, eutrophication and degraded water quality are becoming major threats to these important habitats [18, 21, 25] with seagrass coverage declining at a rate of 110 km^2^ yr^-1^ worldwide since 1980 [26]. Due to their ecological importance seagrasses themselves have been well studied, however, studies on the microbial communities (bacteria and archaea) that occur on the surfaces of seagrass blades are limited [27–30] and their role as a part of the seagrass holobiont is poorly understood.

This study characterizes the microbial communities found within the phyllosphere of a tropical seagrass, *Thalassia testudinum* Banks ex Kӧnig (turtle grass), and examines the relative influence of the plant-host and water column conditions on microbial community composition. To our knowledge, this study contains the most in-depth sampling of the microbial communities on seagrass blades to date and is the first to do so in conjunction with seagrass surveys and environmental sampling. The gradient of environmental conditions due to riverine input at these sampling sites offers an ideal natural setting to examine the influence of relatively small environmental changes on phyllosphere microbial community structure within the larger context of bay-wide environmental conditions. From past studies [16, 29, 30], we expect the phyllosphere microbial communities on *T. testudinum* to differ significantly from the free-living communities of the water column. However, depending on the influence of plant-host and environmental conditions, the variation between microbial communities can be expected to occur at different scales. Therefore, we tested the following two hypotheses (H). H1: If the environmental conditions in the surrounding water column are the major determinant, we would expect to see that microbial compositional changes are correlated with differences in environmental conditions, which can occur across sites and sampling date. H2: Environmental conditions can alter seagrass host characteristics and the host plant’s condition. For instance, water temperature, depth (i.e. light availability), and salinity are all known to affect *T. testudinum* growth, and outside of the optimum range, can act as stressors for the plant [19, 31, 32]. In this case, differences in host plant condition and characteristics among sites, but not sampling date, will influence the identity and composition of blade surface microbial communities.

## Materials and Methods

### Sampling Location

The study sites were located in Apalachee Bay, Florida in the northeastern Gulf of Mexico. The coast of this area is mostly undeveloped with the St. Marks National Wildlife Refuge occupying the adjacent land. Five sites (ABT-1 – ABT-5) were established starting near the mouth of the St. Marks River (30.07059°N, 84.16687°W) and extending south in a linear fashion approximately three kilometers into the bay (30.04194°N, 84.16634°W; Fig 1). These sites occur across a gradient of environmental conditions caused by riverine input with the farthest site from shore located on a shoal. Seagrasses in this region form dense meadows with mixed species composition. As the dominant species at all study sites, *Thalassia testudinum* was the focus of this study, however less abundant species were also present, including *Syringodium filiforme* Kützing (manatee grass), *Halodule wrightii* Ascherson (shoal grass) and *Halophila engelmannii* Ascherson (star grass).

**Fig 1.**
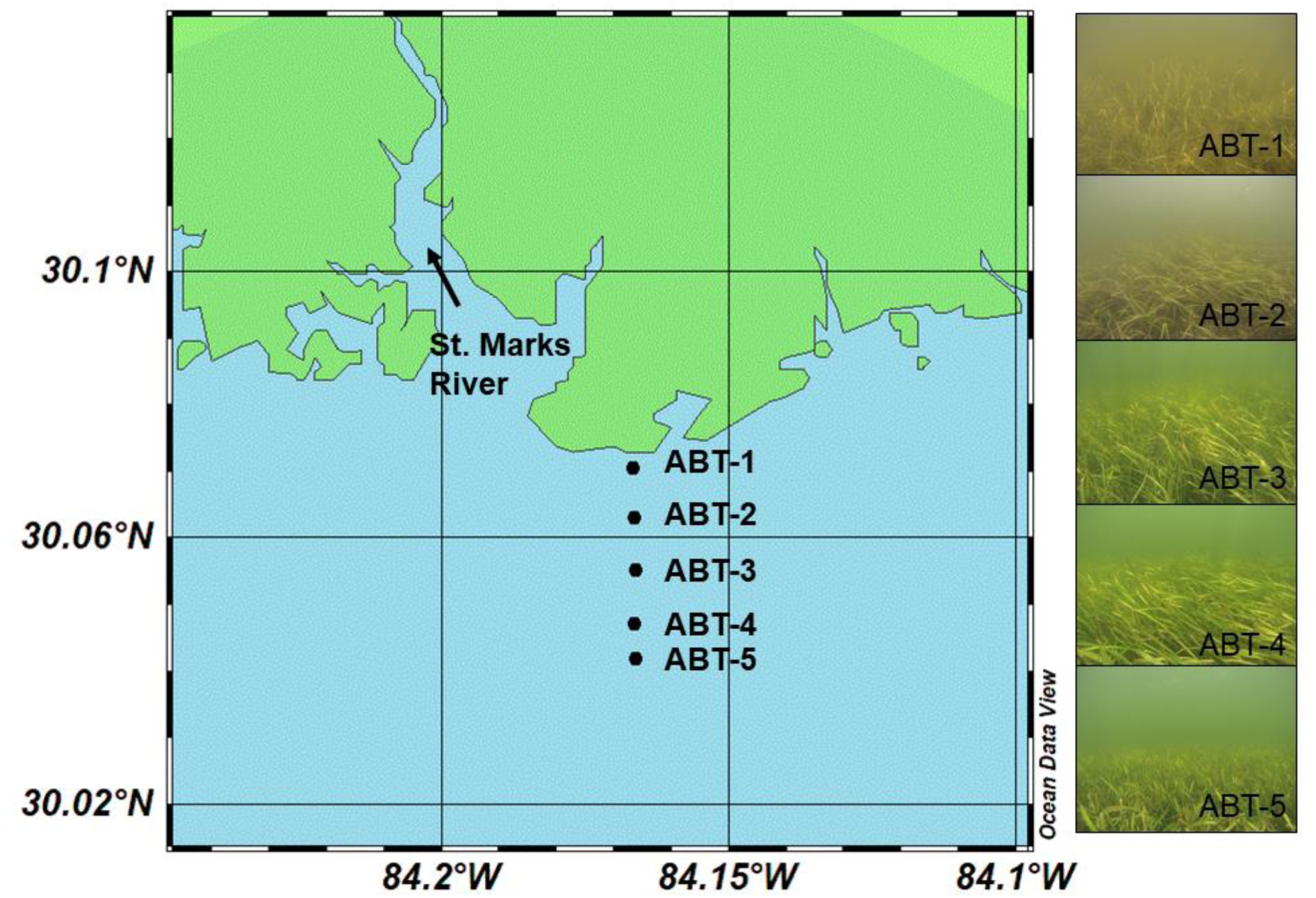
Map (left) showing orientation of five study sites (ABT-1 to ABT-5) in relation to the coastline and St. Marks River mouth. Right panel shows conditions at the five study sites taken on July 22, 2016. Map created with Ocean Data View [33].

### Microbial Sampling

To determine microbial community structure and diversity, samples were taken at all five sites (Fig 1) on three separate dates (July 22, August 20, and September 21, 2016) capturing both spatial and temporal variation. On each date, samples were taken from both the blade surface and water column with all sites visited during a six-hour period. To sample water column communities, one liter of seawater was collected from above the seagrass canopy, filtered using a sterile syringe with a 2.7 µM pre-filter, and microbial biomass was collected on a 0.22 μM Sterivex™ filter. To sample the blade surface microbial communities, five *T. testudinum* shoots were chosen randomly at each site at least one meter apart. One blade from each shoot was then removed using sterile, disposable forceps and a sterile swab (PurFlock® Ultra, Puritan Diagnostics, LLC) was used to sample the blade surface microbial communities. Microbial sampling was standardized by using the second oldest blade in the shoot and only swabbing healthy tissue free of algal epiphytes. Microbial samples (swabs and Sterivex™ filters) were immediately fixed in RNAlater® and placed on dry ice to be transported to Florida State University where all samples were stored at −80°C until further processing.

### Total Suspended Solid and Nutrient Analyses

On all sampling dates, 0.22 μM filtered seawater was also collected at each site, stored in acid-washed 30 ml Nalgene™ bottles, and placed on ice for nutrient analysis. These samples were kept frozen and sent to the Marine Chemistry Laboratory at the University of Washington, School of Oceanography for analysis of PO 3-, NO -, NO, and NH following the protocols of the WOCE Hydrographic Program. For total suspended solids (TSS), 1 liter of unfiltered seawater was collected at each site into acid-washed 500 ml HDPE bottles and placed on ice. These water samples were immediately processed on return to the laboratories at Florida State University by filtering through a pre-weighed 0.7 μM filter using a vacuum pump. Filters were then dried at 60°C for 48 hours and re-weighed to determine TSS content.

Additional *T. testudinum* blades were randomly chosen from a 20 m^2^ area at each site and sampling date for nutrient analysis. Immediately after sample collection*, T.* testudinum blades were cleaned of epiphytes and debris by hand and use of razor blade and then dried at 60°C for 72 hours in the laboratory at Florida State University. Dried samples were then ground to a fine powder using an acid-washed mortar and pestle and stored in glass screw top vials. These samples were sent to the Stable Isotope Ecology Laboratory of the Center for Applied Isotope Studies at the University of Georgia, where they were analyzed for total carbon, nitrogen, and phosphorus content, as well as stable carbon (δ^13^C) and nitrogen (δ^15^N) isotope ratios for indication of short-term and long-term nutrient conditions at each site. Environmental parameters, including water temperature, salinity, conductivity, and dissolved oxygen, were also measured at all sites using a YSI Pro2030.

### Seagrass Surveys

Simultaneous with the microbial sampling, seagrass surveys were conducted to characterize seagrass host condition. At each site, a permanent transect was established running east to west with 1 m^2^ quadrats 15 meters apart along the transect to determine seagrass abundance (percent cover). Within each quadrat, ten blades were randomly chosen to be measured for morphometrics (blade width and length) with only the second and third oldest blades used. Leaf growth rates were measured as an indicator of seagrass growth and leaf turnover. Sexual reproduction was not quantified as clonal reproduction is thought to be the dominant form of growth form in the northern Gulf of Mexico [34] and as only one inflorescence was observed during the study period. Growth rates were measured using a modification of the leaf marking method originally described by Zieman [35] in which shoots were randomly chosen at each site and marked two centimeters above the leaf sheath with a 1/16-inch hand punch. Marked shoots were collected after three weeks and new material was measured by length and dry weight. Water depth, temperature, and salinity were also measured at each site during these surveys to further capture the variation in environmental conditions over the course of the study period.

### Microbial Community Analysis

DNA was extracted from the blade surface and water column microbial samples using a phenol-chloroform extraction method [36] and then purified using the QIAGEN AllPrep™ DNA/RNA Mini Kit. 16S rRNA genes were amplified from DNA extracts in duplicate in accordance with the protocol described by Caporaso et al. [37, 38] using a modified annealing temperature of 60°C with the archaeal and bacterial primers 515F and 806R (targets the V4 region of *E. coli*) modified by Apprill et al. [39] and Parada et al. [40]. During this stage, some samples from 2016 did not successfully amplify and were excluded from sequencing. Amplicons were sequenced using an Illumina MiSeq in 250 x 250 b.p. mode. These sequences will be available in NCBI’s SRA database (BioProject PRJNA623164) and on the Mason server at http://mason.eoas.fsu.edu. Raw sequences were demultiplexed using QIIME2 v. 2019.7 [41]. Demultiplexed reads were quality filtered, including chimera removal, and joined using DADA2 [42]. The resulting amplicon sequence variant (ASV) table was filtered to remove any sequences resulting from mitochondrial or chloroplast DNA and normalized using cumulative sum scaling (CSS) [43]. Taxonomy was assigned using the SILVA v. 132 [44] database in QIIME2. Alpha diversity metrics, including richness (chao1) and diversity (Shannon-Weiner), were determined after multiple rarefactions were performed on the data to account for differences in sequencing depth in QIIME2.

Statistical analyses were performed using R v 3.6.1 [45] and the vegan package [46] to obtain measures of variation in community structure among sample types, sampling dates, and sites on CSS normalized ASV data. Microbial community dissimilarity was determined using Non-metric Multidimensional Scaling (NMDS) ordination analysis with Bray-Curtis distance using CSS normalized ASV counts. Ordination analyses were performed using the *metaMDS* command with the vegan package [46] in R with 999 permutations and appropriate number of axes to minimize stress. PERMANOVA (*adonis*) from the vegan package [46] in R was used to determine significant differences in community composition among sample types, sites, and sampling dates with CSS normalized ASV counts as the response variable. Environmental fitting (*envfit*) from the vegan package [46] in R was also used with the resulting ordinations to show correlations between environmental/host conditions and changes in community member identity and abundances. As the distribution of CSS normalized ASV data was determined to not be normal using the Shapiro-Wilkes test, non-parametric statistical tests were used including Dunn’s test and Wilcoxon rank sum test to determine significant differences in alpha diversity among sites and sampling dates. Non-parametric tests were also used to determine differences in environmental and host characteristics among sites and sampling dates. During the PCR step, some of the blade surface samples (n=23) failed to amplify and could not be sequenced. Due to this and uneven sample sizes between blade surface and water samples, significant differences were confirmed by subsampling down to the minimum sample size using bootstrapping without replacement for 1000 permutations. Non-parametric tests were then performed on bootstrapped data to determine significant differences in richness and diversity. Mean values are reported with plus or minus the standard error.

## Results

### Host Plant and Site Characterization

Despite the sites being relatively close in proximity (< 3 km apart), differences were observed in environmental conditions across site and over the three sampling dates. On average, salinity increased with distance from shore ranging from 21 ppt at the first site (ABT-1) to 27 ppt at the fifth site (ABT-5; Table 1). Differences in salinity among sites were only marginally significant (Kruskal-Wallis, p=0.05); however, salinity at all sites was significantly different among sampling dates (Kruskal-Wallis, p<0.01) as the July sampling had a significantly higher average salinity (29.2 ppt) than the August sampling (19.7 ppt; Dunn’s test with Bonferroni correction, p<0.01). Water depth was significantly different by site (Kruskal-Wallis, p<0.01), however due to the bathymetry of the area, water depth did not co-vary with distance to shore as might be expected (Fig 1). Water depths at the farthest site from shore (ABT-5) had an average depth (1.2 m) similar to the closest site (ABT-1; 0.98 m; Table 1). Additionally, water depths at the deeper, intermediate sites were similar ranging on average from 1.6 m at ABT-2 to 2.8 meters at ABT-4. The fourth site (ABT-4) was found to have a significantly greater depth (2.18 m) than the first site (ABT-1; Dunn’s test with Bonferroni correction, p<0.01) and ABT-3 was found to have a marginally greater depth than the first site (ABT-1) and ABT-4 had a marginally greater depth than the farthest site (ABT-5; Dunn’s test with Bonferroni Correction, p=0.09 and p=0.07, respectively). Dissolved oxygen concentrations (DO; mg/l), total suspended solid content (TSS; mg/l), and water column nutrients (PO_4_^3-^, NO_3_^-^, NO_2_^−^ and NH_4_^+^) were not significantly different by site or sampling date (Kruskal-Wallis, p<0.05; Table 1).

**Table 1.** Site Characterization. Mean water column and host parameters (± S.E.) measured at each site during seagrass surveys and microbial samplings during 2016.

| Water Column Parameters |  |  |  |  |  |  |  |  |  |  |
| --- | --- | --- | --- | --- | --- | --- | --- | --- | --- | --- |
|  | Depth | Temp. | Salinity (‰) | DO (mg/l) | TSS | PO4 | NO3 | NO2 | NH4 |  |
|  | (m) | (°C) |  |  | (mg/l) | (µg/l) | (µg/l) | (µg/l) | (µg/l) |  |
| ABT-1 | 0.98 ± | 30.7 ± | 21.0 ± 1.8 | 4.33 ± | 15.88 ± | 1.9 ± | 4.1 ± | 1.7 ± | 56.8 ± |  |
|  | 0.08 | 0.8 |  | 1.07 | 4.98 | 0.5 | 2.6 | 0.6 | 28.0 |  |
| ABT-2 | 1.6 ± | 30.6 ± | 22.7 ± 1.8 | 4.98 ± | 9.77 ± | 2.0 ± | 0.1 ± | 1.7 ± | 34.7 ± |  |
|  | 0.20 | 0.4 |  | 0.57 | 1.60 | 0.1 | 0.1 | 0.8 | 6.2 |  |
| ABT-3 | 1.85 ± | 30.8 ± | 24.9 ± 2.2 | 6.01 ± | 9.62 ± | 1.9 ± | 1.3 ± | 2.2 ± | 121.9 |  |
|  | 0.15 | 0.1 |  | 0.04 | 1.65 | 0.5 | 0.6 | 1.0 | ± 50.0 |  |
| ABT-4 | 2.18 ± | 30.8 ± | 26.0 ± 2.0 | 5.60 ± | 11.53 ± | 1.1 ± | 0.8 ± | 0.6 ± | 8.8 ± |  |
|  | 0.13 | 0.2 |  | 0.42 | 3.07 | 0.1 | 0.5 | 0.1 | 0.7 |  |
| ABT-5 | 1.2 ± | 30.8 ± | 27.3 ± 2.0 | 5.55 ± | 19.76 ± | 1.3 ± | 0.1 ± | 0.9 ± | 13.0 ± |  |
|  | 0.03 | 0.4 |  | 0.43 | 2.47 | 0.3 | 0.1 | 0.6 | 7.1 |  |
| Seagrass Host Parameters |  |  |  |  |  |  |  |  |  |  |
|  | Length | Width | GR | GR | Cover | TP (%) | TN | TC (%) | δ<sup>15</sup>N | δ<sup>13</sup>C |
|  | (cm) | (cm) | (mg/s/day) | (cm/s/day) | (%) |  | (%) |  | (‰) | (‰) |
| ABT-1 | 29.9 ± | 0.4 ± 0.0 | 1.57 ± 0.27 | 0.9 ± 0.2 | 46 ± 12 | 0.146 | 2.24 ± | 33.93 | 1.26 ± | -16.12 |
|  | 0.8 |  |  |  |  | ± 0.01 | 0.08 | ± 0.28 | 0.32 | ± 0.73 |
| ABT-2 | 31.3 ± | 0.7 ± 0.1 | 4.31 ± 0.26 | 1.3 ± 0.1 | 66 ± 5 | 0.133 | 2.11 ± | 33.66 | 2.22 ± | -12.79 |
|  | 0.8 |  |  |  |  | ± 0.00 | 0.01 | ± 0.14 | 0.13 | ± 0.75 |
| ABT-3 | 43.6 ± | 0.7 ± 0.0 | 5.31 ± 0.57 | 1.8 ± 0.1 | 73 ± 10 | 0.136 | 2.14 ± | 33.52 | 1.63 ± | -12.76 |
|  | 1.2 |  |  |  |  | ± 0.01 | 0.05 | ± 0.27 | 0.36 | ± 0.77 |
| ABT-4 | 40.0 ± | 0.7 ± 0.0 | 5.38 ± 0.39 | 1.6 ± 0.1 | 55 ± 11 | 0.130 | 2.02 ± | 33.52 | 3.00 ± | -11.93 |
|  | 1.3 |  |  |  |  | ± 0.00 | 0.11 | ± 0.28 | 0.12 | ± 0.82 |
| ABT-5 | 21.5 ± | 0.7 ± 0.0 | 2.05 ± 0.15 | 0.7 ± 0.0 | 86 ± 12 | 0.138 | 2.11 ± | 36.55 | 3.02 ± | -12.50 |
|  | 0.4 |  |  |  |  | ± 0.01 | 0.14 | ± 2.96 | 0.04 | ± 0.79 |

*Thalassia testudinum* growth morphology also varied among the five sites with seagrass characteristics the most similar at intermediate sites. Blade lengths had a significant positive correlation with water depth (p<0.01, adjusted R^2^=0.51) and were greatest at the intermediate sites (ABT-2, ABT-3, and ABT-4; Table 1). Blade lengths were found to be significantly different at all sites with the exception of the third and fourth sites (Dunn’s test with Bonferroni correction, p<0.01). However, blade widths did not follow this same trend with depth as the widths at the first site (ABT-1) were found to be significantly smaller than at all other sites, including the fifth site of similar depth (Dunn’s test with Bonferroni correction, p<0.001). Leaf growth rates were found to co-vary with blade width and lengths and were highest at sites ABT-4 and ABT-3 (5.38 and 5.31 mg/day/shoot, respectively), which also had the longest blades on average (41.56 cm and 39.49 cm, respectively) and greatest average water depth (Table 1). Leaf growth rates (mg/day/shoot) were found to be significantly higher at ABT-3 than at both the closest and farthest sites (ABT-1 and ABT-5) and significantly higher at ABT-4 than at the closest site (ABT-1; Dunn’s test with Bonferroni correction, p<0.05; Table 1). Blade nutrient composition (%N, %C, %P, and δ^13^C content) was not found to be significantly different due to site or sampling date; however mean δ^15^N was found to have significantly higher concentrations at the farthest site (ABT-5) than the first site (ABT-1; Dunn’s test with Bonferroni correction, p<0.05; Table 1). Additionally, percent cover of *T. testudinum* was found to be significantly greater at the farthest site (ABT-5; 86%) than at the first (ABT-1; 46%) and the fourth site (ABT-4; 55%; Dunn’s test with Bonferroni correction, p<0.05; Table 1).

### Blade Surface Community Composition and Diversity

A total of 11,252 ASVs were found in *T. testudinum* blade surface samples (n=52) across all sites and sampling dates. Samples were dominated by bacteria, specifically the phyla Proteobacteria, Cyanobacteria, and Planctomycetes, with archaea making up an average of 0.25% of blade surface community relative abundance. While at the ASV level there were no dominant taxa, two bacterial classes, Gammaproteobacteria (21.56 ± 4.78%) and Alphaproteobacteria, (20.26 ± 6.94%) each comprised approximately 20% of community relative abundance on average (CSS normalized data was converted to relative proportions). The majority of microorganisms (ASVs) comprised less than 1% of sample relative abundance, however three ASVs belonging to the cyanobacterial family Cyanobiaceae were the exception, including the cultured bacterium *Synechococcus* sp. CENA143. These ASVs comprised a combined 5% of the community relative abundance on average. These three abundant cyanobacteria are also closely related to known cyanobacteria found in mangrove systems [47, 48], including *Synechococcus* sp. CENA 172 and CENA 180 (100% similarity, GenBank Accession KC695872.1 and KC695865.1).

On average, blade surface communities had a richness (chao1) of 792.12 ± 190.92 ASVs with the maximum richness (1565.75 ASVs) occurring at the second site (ABT-2) from shore and the lowest richness (436.03 ASVs) occurring at the fifth site (ABT-5) furthest from shore (Fig 2A). Across all sampling dates, richness was significantly lower at the fifth site (657.81 ± 118.99 ASVs) than at the second (873.76 ± 236.06 ASVs) and third sites (895.55 ± 165.01 ASVs; Dunn’s Test with Bonferroni Correction, p<0.05; Fig 2A). Diversity (Shannon-Weiner) showed a similar pattern, with ABT-5 having significantly lower diversity (8.39 ± 0.29) than ABT-2 (8.80 ± 0.30) and ABT-3 (8.86 ± 0.28; Dunn’s Test with Bonferroni Correction, p<0.05; Fig 2B). Both richness and diversity were significantly different by site location, however neither metric significantly differed due to sampling date. Differences in community composition were visualized with NMDS ordination analysis (Fig 3A) with each blade surface community represented by a single point. Blade surface samples were found to differ significantly in species composition due to site and sampling date as well as having a significant interaction between site and sampling date (PERMANOVA, p<0.001). All sites were found to be significantly different within a sampling date and sampling dates were found to be significantly different within each site with the exception of ABT-1 (PERMANOVA, p<0.001).

**Fig 2.**
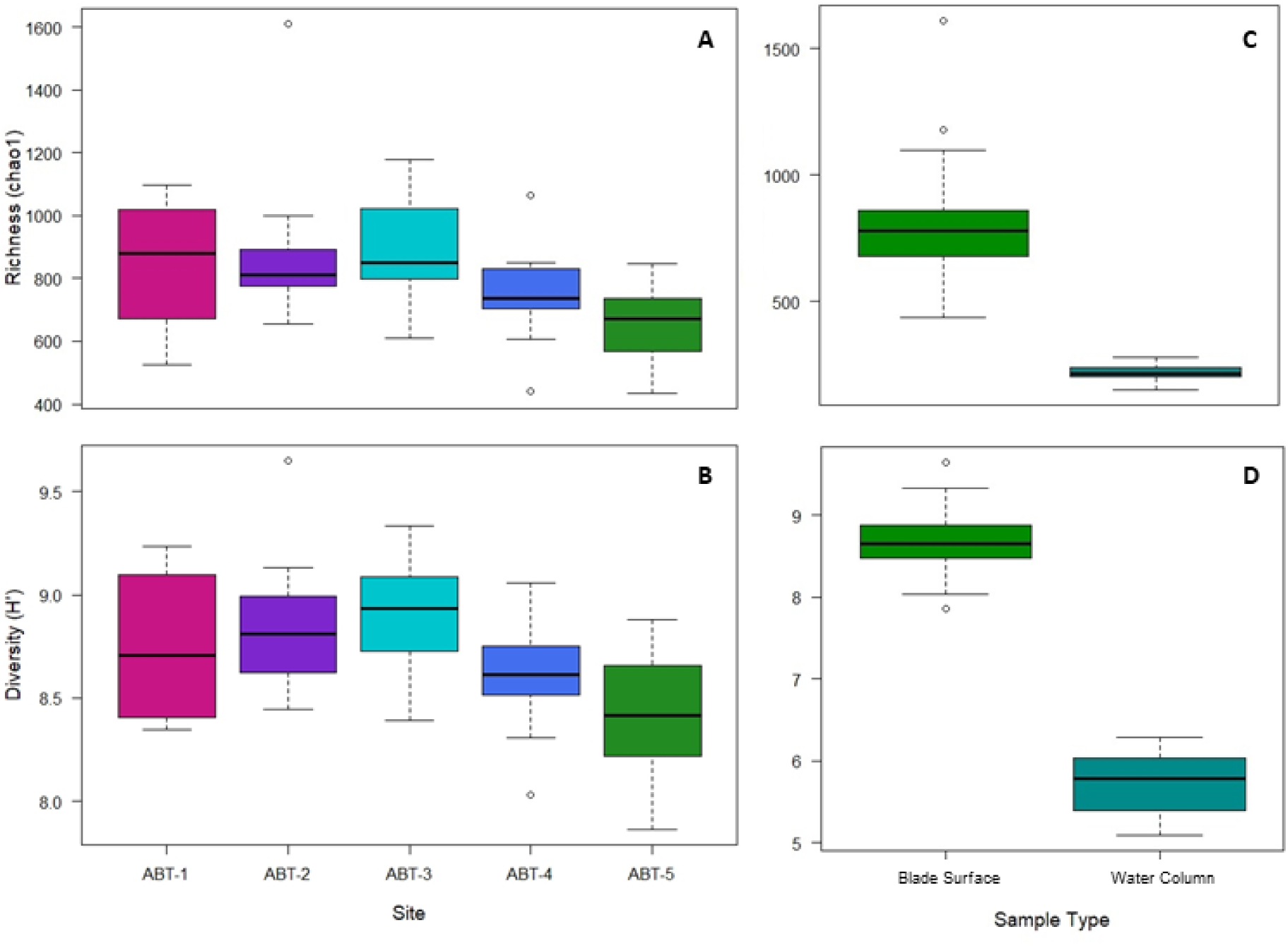
Boxplots showing (A) richness (chao1) and (B) diversity (Shannon-Weiner) of blade surface samples among the five sites from all three sampling dates and differences in (C) richness (chao1) and (D) diversity (Shannon-Weiner) between blade surface and water column samples.

**Fig 3.**
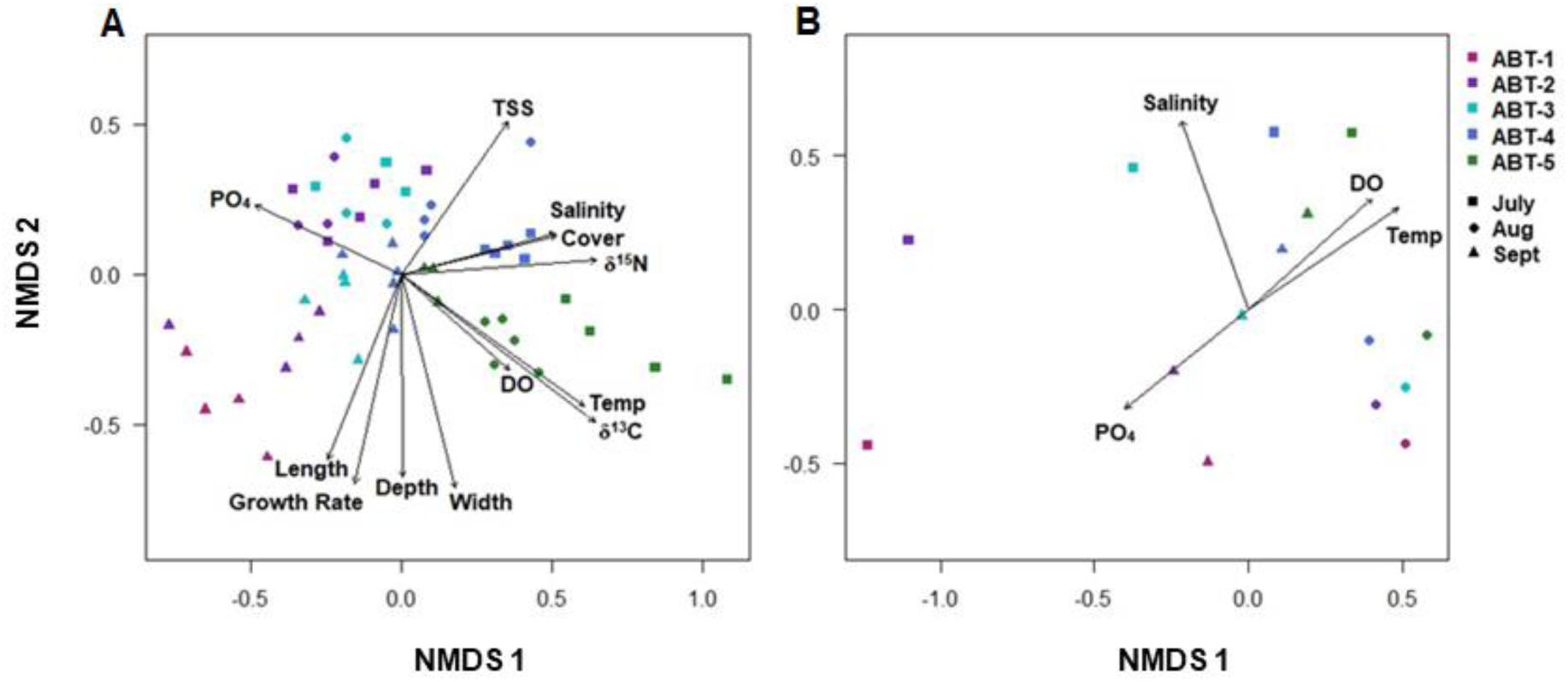
(A) Non-metric Multi-dimensional Scaling (NMDS) Ordination of blade surface 16S rRNA iTag sequence data (stress=0.16, k=2). *Thalassia testudinum* blade surface microbial communities represented as single points and show aggregation by both site and sampling date. Arrows show significant correlations with environmental (water temperature, depth, salinity, total suspended solids content, and phosphate concentration) and host (blade length and width, leaf growth rates, percent cover, blade δ^15^N and δ^13^C content) characteristics (*envfit*, corrected p<0.05). Direction of arrow represents correlation with changes in community composition and length of arrow represents R^2^ values. (B) Non-metric Multi-dimensional Scaling (NMDS) Ordination of free-living water column 16S rRNA iTag sequence data (stress=0.06, k=2). Water column communities show significant differences by site. Arrows show environmental parameters (salinity, water temperature, dissolved oxygen content, and phosphate concentration) that were found to significantly correlate with community composition (*envfit*, corrected p<0.05). For A and B, symbol shape represents site (ABT-1: pink, ABT-2: purple; ABT-3-light blue, ABT-4: dark blue, ABT-5: green) and shape represents sampling date (July: square, August: circle, September: triangle).

### Correlations with Environmental and Host Characteristics

Differences in microbial community composition among the blade surface samples were found to have significant correlations with environmental factors at each site, including water temperature, average depth, salinity, total suspended solids concentration, and phosphate concentration as well as host characteristics, including average blade length, average blade width, average percent cover, *T. testudinum* leaf growth rates, and stable isotope composition (δ^15^N and δ^13^C content) (Fig 3A; Table 2). In addition, dissolved oxygen content in the water column had a marginally significant correlation with blade surface composition (Table 2).

**Table 2.** Environmental fitting results. Environmental fitting (*envfit*, vegan package) on the NMDS ordination was used to determine significant environmental and host characteristics (p<0.05, indicated by asterisk). Bonferroni correction was applied to p-values to account for multiple comparisons.

|  | NMDS1 | NMDS2 | R<sup>2</sup> | p-value | corrected p-value |
| --- | --- | --- | --- | --- | --- |
| **<u>Host Characteristics</u>** |  |  |  |  |  |
| Avg. Length (cm) | -0.37028 | -0.92892 | 0.3974 | 0.001 | 0.018 * |
| Avg. Width (cm) | 0.24096 | -0.97054 | 0.4877 | 0.001 | 0.018 * |
| Growth Rate (g/shoot/day) | -0.21823 | -0.9759 | 0.4596 | 0.001 | 0.018 * |
| Avg. Cover (%) | 0.96926 | 0.24605 | 0.2523 | 0.002 | 0.036 * |
| Total N (%) | -0.99989 | -0.01481 | 0.0636 | 0.200 | 1.000 |
| Total C (%) | 0.78755 | 0.61625 | 0.1329 | 0.025 | 0.450 |
| Total P (%) | -0.89918 | 0.43758 | 0.0789 | 0.136 | 1.000 |
| δ<sup>15</sup>N (‰) | 0.99711 | 0.07593 | 0.3810 | 0.001 | 0.018 * |
| δ<sup>13</sup>C (‰) | 0.79302 | -0.60919 | 0.5883 | 0.001 | 0.018 * |
| **<u>Abiotic Characteristics</u>** |  |  |  |  |  |
| Water Temp. (°C) | 0.81146 | -0.58440 | 0.5076 | 0.001 | 0.018 * |
| Avg. Depth (m) | 0.00464 | -0.99999 | 0.4144 | 0.001 | 0.018 * |
| Dissolved Oxygen (mg/l) | 0.74554 | -0.66646 | 0.2075 | 0.003 | 0.054 |
| Salinity (‰) | 0.96511 | 0.26185 | 0.2539 | 0.001 | 0.018 * |
| Total Suspended Solids (mg/l) | 0.56713 | 0.82363 | 0.3463 | 0.001 | 0.018 * |
| Phosphate (µg/l) | -0.90468 | 0.42609 | 0.2643 | 0.002 | 0.036 * |
| Nitrate (µg/l) | -0.44827 | -0.89390 | 0.1733 | 0.012 | 0.216 |
| Nitrite (µg/l) | -0.71556 | -0.69855 | 0.1008 | 0.059 | 1.000 |
| Ammonium (µg/l) | -0.92074 | -0.39018 | 0.0965 | 0.079 | 1.000 |

In addition, other community metrics of the blade surface samples, including diversity and richness, correlated with both environmental and host parameters. Shannon diversity of the blade surface communities was significantly correlated with total suspended solids concentration, phosphate concentration, and blade δ^15^N content (Spearman’s Rank Order Correlation with Bonferroni correction, p≤0.05). Richness (chao1) of the blade surface communities was also significantly correlated with water column total suspended solids and blade δ^15^N content (Spearman’s Rank Order Correlation with Bonferroni correction, p<0.05). However, most of these correlations with environmental and host characteristics are confounded as parameters. such as water depth and blade lengths, co-vary.

### Blade Surface vs. Water Column Microbial Communities

In evaluating free-living microbial communities in the water column samples (n=15) the number of total ASVs (1,083) was lower than in blade surface samples; however, approximately 80% of those ASVs were also present in blade surface communities. Individual samples had significantly lower richness (215.92 ± 36.97 ASVs; Fig 2C) and diversity (5.73 ± 0.40; Fig 2D) than the blade communities (Wilcoxon rank sum test, p<0.001). Community composition of water column samples was found to be significantly different from the composition of blade surface samples (PERMANOVA, p=0.001). Within water column samples, community composition differed significantly by site (PERMANOVA, p<0.05; Fig 3B), but not by sampling date.

The composition of the free-living water column microbial communities was significantly correlated with several environmental parameters, including water temperature, salinity, dissolved oxygen, and phosphate concentration (*envfit* with Bonferroni correction, p<0.05; Fig 3B). Salinity and water temperature explained the most variance with the highest R^2^ values of 0.93 and 0.75, respectively. However, in contrast to blade surface communities, diversity and richness of the water column communities did not significantly correlate with any of the environmental conditions (Kruskal-Wallis, p>0.05).

### Core Community

Although blade surface communities contained many species in low abundances, 21 ASVs were present in 100% of the blade surface samples and combined comprised 8-21% of relative abundance (Fig 4). These 21 ASVs represent the following five different bacterial phyla in order of decreasing average relative abundance-Cyanobacteria, Proteobacteria, Planctomycetes, Chloroflexi, and Bacteroidetes (Fig 4). The three most abundant ASVs from the blade surface samples, which were discussed above, were also amongst these 21 core community members. The combined abundance of these 21 ASVs was also found to have a significant positive correlation with water temperature at the time of sampling (R^2^=0.49), average water depth (R^2^=0.34), and leaf growth rates (R^2^=0.34; Spearman’s Rank Order Correlation with Bonferroni correction, p<0.05). Additionally, combined abundances were found to be significantly different among sites (Kruskal-Wallis, p<0.001) with the furthest two sites (ABT-4 and ABT-5) having a significantly higher abundance of the core community than the first two sites (ABT-1 and ABT-2; Dunn’s test with Bonferroni correction, p<0.01). Members of the core community were largely absent from the water column community. The only exceptions were two ASVs that occurred in no more than two water samples and in extremely low abundances (<0.006% on average).

**Fig 4.**
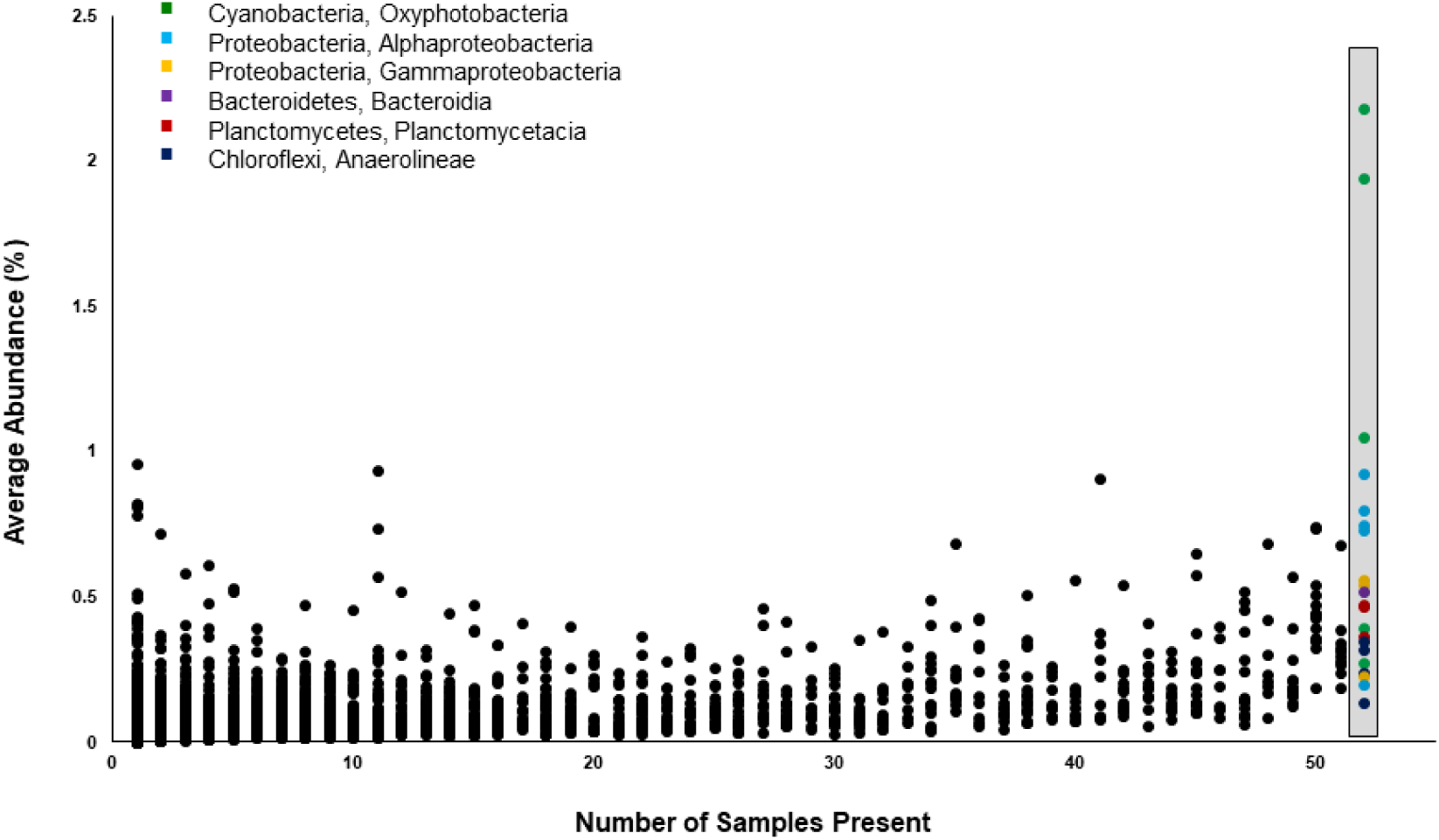
The average relative abundance of each of the 11, 252 ASVs found in blade surface samples is plotted against the number of samples that included that ASV. Twenty-one ASVs were found in 100% of blade surface samples (n=52) and comprise a potential ‘core’ community for the *T. testudinum* phyllosphere. Core community members, highlighted in the gray box, consisted of both abundant and rare species. The core community members are color coded by bacterial phylum and class (green-Cyanobacteria, Oxyphotobacteria; light blue-Proteobacteria, Alphaproteobacteria; yellow-Proteobacteria, Gammaproteobacteria; purple-Bacteriodetes, Bacteroidia; red-Planctomycetes, Planctomycetacia; dark blue-Chloroflexi, Anaerolineae).

## Discussion

### Variation within the Phyllosphere Microbial Communities

This study uses the scale and pattern of variation among *T. testudinum* blade surface microbial communities to investigate the relative influence of environmental conditions (H1) and host characteristics (H2) in determining community structure. Results showed that blade surface microbial community composition, which includes both ASV identity and abundance, varied significantly among the five sites and sampling dates (July, August, and September) revealing that *T. testudinum* blades can host a diversity of microbes and that these blade surface microbial communities can be highly variable over a relatively short distance (<3 km) and time period (3 months). These results are consistent with the hypothesis that environmental conditions can have a pronounced influence on these blade surface microbial communities (H1). Salinity and water depth showed the most marked differences among sampling date and site, respectively, and could be driving the compositional changes in blade surface microbial communities that were observed among sampling date and site. Interestingly, differences in microbial community composition correlated with additional environmental variables aside from salinity and water depth suggesting that small changes in environmental conditions can have a large influence on blade surface microbial community structure. However, some of these parameters co-vary and it cannot be determined from these results alone which environmental factors are driving these compositional changes.

Additionally, seagrass surveys revealed differences in *T. testudinum* plant characteristics (blade length, blade width, and blade δ^15^N content) among sites and these differences in the host plant characteristics could also affect microbial composition among sites. Correlations between these site differences in seagrass characteristics as well as leaf growth rates, percent cover, and δ^13^C content and microbial community composition may indicate that condition of the host plant influences the blade surface communities (H2). However, this study cannot distinguish the direct effects of the environment (H1) or the effects of the environment through host plant traits (H2) as the cause for the observed changes in composition. For instance, low salinity has been shown to result in thinner *T. testudinum* blade widths [49] and, since microbial community composition correlated significantly with both factors, it is not known if these differences are a result of the environment or the host’s condition. Manipulative studies are needed to separate the relative influences of host plant traits and environment conditions on microbial community structure.

### A Possible ‘Core’ Community

While the blade surface microbial communities were highly variable across sampling dates and sites, 21 ASVs were found in all *T. testudinum* blade surface samples. These 21 microorganisms may comprise a ‘core’ community that is possibly unique to *Thalassia* blades. While a few members are abundantly dominant ASVs, many of the 21 ASVs in the core community are rare (Fig 4). The ubiquity of these ASVs may indicate that they are ecologically significant in determining community structure or that they perform key community functions. Members of this core community may perform processes that are beneficial or deleterious to the host and, in turn, could affect the maintenance of a healthy microbial community on these blade surfaces. Core community members may also be important foundation or key stone species for the entire community involved in biofilm production or in determining community structure [50, 51]. Members of the core community found in this study belonged to six bacterial classes (Fig 4), including Gammaproteobacteria, Alphaproteobacteria, and Anaerolinae. Other studies have shown that bacteria from these classes are abundant in the microbiome of *T. testudinum* and other seagrasses [27, 29, 30, 52]. Additional studies need to be conducted to confirm the presence of a *T. testudinum* core community on greater spatial and temporal scales and to determine under what conditions this core community persists [30]. Furthermore, it may be that the core community members are defined by functional group rather than taxonomic identity [53]. Functional analysis, including metatranscriptomics and metaproteomics, could reveal if assembly of core community members is dictated by function rather than taxonomy. Identifying functions encoded in the core and greater blade surface communities will help to understand the relationship these microbial communities have with their seagrass host.

### Comparisons with Water Column Microbial Communities

The composition of the microbial communities associated with *T. testudinum* blade surfaces was found to be highly diverse and, as shown in previous studies the composition of these communities was significantly different from that of the free-living communities in the water column [16, 29, 30]. Water samples also had significantly lower richness and diversity than *T. testudinum* blade surface communities, similar to the findings of Ugarelli et al. [30]. This suggests that there are source populations for blade surface microbial communities other than the surrounding water column, including the sediment and rhizosphere microbial communities. Results from this study differ from a previous study on *T. testudinum* phyllosphere microbial communities [30] in which the number of ASVs found in the phyllosphere (3,347) was much lower. The higher number of ASVs observed in this study (over 11,000) could be due to differences in sampling methods as well as including a larger sample size. In the temperate seagrass *Zostera marina* Linneaus, Crump et al. [29] found that on average 13% of phyllosphere communities were dominated by a single OTU, which differs from this study in which ASVs were used as the taxonomic unit and the highest relative abundance was found to be 2% of the community for a single ASV. However, it could be expected that seagrasses living in temperate climates have different leaf microbiomes than tropical seagrasses due to shorter growing seasons and larger seasonal fluctuations in environmental conditions. Additionally, comparisons across seagrass microbiome studies are not always appropriate due to differences in sampling methods, sequencing platforms, and data pipelines used. This highlights the need for a convergence on methods in order to make larger generalizations about the structure and roles of seagrass phyllosphere microbial communities.

### Comparisons with the Terrestrial Phyllosphere

Despite the large differences in environmental conditions, there are similarities in microbial community structure and composition between the marine and terrestrial phyllospheres. On a broad scale, both community types are dominated by members of the bacterial domain with relatively low abundances of archaea. Further, Proteobacteria (including alpha- and gammaproteobacteria), Chloroflexi, and Bacteroidetes were found to be dominant taxonomic groups in this study as well as in the leaf surfaces of terrestrial plants [54–56]. The results from this study indicate source populations other than the microbial communities of the water column exist, which is similar to the terrestrial phyllosphere where sediment microbial communities are considered a major source population for leaf surface communities [3, 5, 57]. Additionally, environment and host characteristics appear to be important drivers for both marine and terrestrial phyllosphere microbial community structure. Abiotic factors, including precipitation/moisture, temperature, radiation, and pollution [3, 6–8, 10], as well as biotic factors, such as plant-host identity, leaf age, plant growth, and plant functional traits [9, 56, 58] have been found to correlate with microbial community composition on terrestrial leaf surfaces.

Core microbial communities, which are theorized to have ecological importance, have also been identified on the leaves of terrestrial plants. In the terrestrial phyllosphere, core community members have been found to comprise a small percentage of relative taxonomic diversity yet can comprise up to 73% of sequence abundance [8, 56]. A similar pattern was found in this study with core community members comprising 0.19% of taxonomic richness, but up to 21% of sequence abundance. While core communities in the terrestrial phyllosphere have been found to contain members of the same phyla, including Proteobacteria and Bacteroidetes, at a finer taxonomic scale, community members differ. The phyllosphere of *T. testudinum* contained several Cyanobacteria as well as members of the family Rhodobacteraceae, which are largely absent from the core communities of terrestrial plants [2, 3, 8, 56–58]. This may indicate that core community members perform different ecological functions in these two environments. For instance, members of the family Rhodobacteraceae have been found to be key community members early on in biofilm formation [59–61], which may be important for community development within the marine phyllosphere.

## Conclusions

This study shows that *T. testudinum* phyllosphere hosts rich microbial communities that are distinct from that of the water column. Further, these blade surface communities can exhibit high variability among sites within a single water basin. Environmental conditions are the most likely driver of differences in blade surface microbial community composition. However, host plant characteristics may also influence the composition of blade surface microbial communities, similar to what is observed in the terrestrial phyllosphere. This correlative study provides the foundation for experiments to more directly investigate the relationship between *T. testudinum* health and the blade surface microbiome. Understanding the role these blade surface communities have as a part of the seagrass holobiont is imperative to understanding seagrass health as a whole and the importance of microbiomes in maintaining healthy coastal ecosystems. Future studies will have important implications for seagrass conservation and management as the continued loss of seagrass will inevitably lead to greater ecological and economic losses [26].

## Acknowledgements

This study was conducted under Florida Fish and Wildlife Commission permit number SAL-16-1799C-SR.

